# DMENet: Diabetic Macular Edema Diagnosis using Hierarchical Ensemble of CNN’s

**DOI:** 10.1101/712240

**Authors:** Rajeev Kumar Singh, Rohan Gorantla

## Abstract

Diabetic Macular Edema (DME) is an advanced stage of Diabetic Retinopathy (DR) and can lead to permanent vision loss. Currently, it affects 26.7 million people globally and on account of such huge number of DME cases and limited number of ophthalmologists, it is desirable to automate the diagnosis process. Computer-assisted, deep learning based diagnosis will help in early detection, following which appropriate medication can help to mitigate the vision loss. Method: In order to automate the screening of DME, we propose a novel DMENet Algorithm which is built on the pillars of Convolutional Neural Networks (CNN’s). DMENet analyses the preprocessed color fundus images and passes it through a two-stage pipeline. The first stage detects the presence or absence of DME whereas the second stage takes the positive cases and grades the images based on severity. In both the stages, we use a novel Hierarchical Ensemble of CNN’s (HE-CNN). This paper uses two of the popular publicly available datasets IDRiD and MESSIDOR for classification. Preprocessing on the images is performed using morphological opening, Gaussian kernel and the dataset is augmented to solve the class imbalance problem. Results: The proposed methodology achieved an Accuracy of 96.12%, Sensitivity of 96.32%, Specificity of 95.84%, and F–1 score of 0.9609. Conclusion: These excellent results establishes the validity of the proposed methodology for use in DME screening and solidifies the applicability of HE-CNN classification technique in the domain of bio-medical imaging.

## Introduction

Diabetic macular edema (DME) is a complication of Diabetic Retinopathy (DR), and it usually occurs when vessels in the central part of the macula are affected by the fluid accretion [1]. DME is caused due to diabetes which is a chronic disease induced by inherited and/or acquired deficiency in the production of insulin by the pancreas. DME is an advanced stage of DR that can lead to irreversible vision loss [2–4].

Diabetes currently affects more than 425 million people worldwide and is expected to affect an estimated 520 million by 2025. It is estimated that 10% of people who suffer from some form of Diabetes are at the risk of DME. DME currently affects around 26.7 million people globally, and the number is expected to rise to around 50 million by 2025 [3, 5]. About 7.7 million Americans have DR and approximately 750,000 are suffering from DME. [4].

Identifying exudates in fundus images is a standard technique for determining DME [6, 7].The nearness of exudates to the macula determines the severity of DME and the probability for DME augments when the exudates are closer to the macula with the risk being maximum when they are inside the macula [8]. Immediate treatment is required if a person is diagnosed with DME to avoid complete vision loss.

The ratio of ophthalmologists to population in developed countries like USA is 1:15,800 and in developing countries like India is 1:25,000 in urban areas and 1:219,000 in rural areas [9, 10]. The limited number of ophthalmologists cannot keep up with the rapidly increasing number of DME patients and this heavily skewed ratio of ophthalmologists with respect to DME patients is also leading to delayed services. Manual evaluation of DME is not adaptable in a large-scale screening scheme, especially in developing countries where there is a shortage of ophthalmologists [11]. During the screening tests, one out of nine people turns out to be positive cases of DME [12]. Another challenging aspect of the healthcare sector in developing nations is to provide the best and timely diagnosis at an affordable cost. In this kind of a scenario we require an automatic disease discovery framework that can reduce cost and workload, as well as solve the shortage of ophthalmologists by limiting the referrals to those cases that require prompt consideration. The reduction of effort and time of ophthalmologists in diagnosis will be pivotal for arresting the growth of DME cases.

Propelled by these promising possibilities, we propose to develop an effective solution using Deep learning techniques to automatically grade the fundus images. Machine learning techniques have powered many aspects of medical investigations and clinical practices. Deep learning is emerging as a leading machine learning tool in computer vision and has started to command significant consideration in the field of medical imaging. Deep learning techniques, in particular, convolutional neural networks, have rapidly gained prominence for analysis of the medical images.

In this paper, we propose a novel algorithm DMENet which automatically analyses the preprocessed color fundus images. Preprocessing on the images is performed using morphological opening, Gaussian kernel and the dataset is augmented to solve the class imbalance problem. After Preprocessing the images, they are passed through a two-stage pipeline. In the first stage, the algorithm detects for the presence/absence of DME and once the presence of DME is confirmed it is passed through the second stage where the image is graded based on the severity. Both these stages are equipped with a state-of-art technique called Hierarchical Ensemble of Convolutional Neural Networks (HE-CNN).

Our proposed DMENet algorithm in this paper is built on a novel classification structure known as HE-CNN which uses the concept of *ensemble learning*. Ensemble learning is a technique which combines the outputs from multiple classifiers to improve the classification performance of the model [13]. This approach is intuitively used in our daily lives, where we seek guidance from multiple experts, weigh and combine their views to make a more informed and optimized decision. In matters of great importance that have financial or medical implications, we often seek a second opinion before making a decision, for example, having several doctors agree on a diagnosis reduces the risk of following the advice of a single doctor whose specific experience may differ significantly from that of others.

## Related Work

The feasibility of manual assessment for the diagnosis of ophthalmic cases has become practically untenable. Automation of at least the first level diagnosis is a clear requirement to improve the efficacy, affordability, and accessibility of our healthcare system. Over the last two decades, many research groups have extensively worked on automating the diagnosis of ophthalmic problems using color fundus images and OCT images. Image acquisition, processing and diagnosis using color fundus images is comparatively faster and in case of developing nations where cost factor plays a vital role, color fundus imaging scores higher than others. Recently the use of data-driven machine learning and deep learning techniques on classical expert labelled image analysis have gained prominence. Convolutional Neural Networks have become the method of choice for automated grade assessment of DME. The techniques used in existing works on automated DME grading on color fundus images can be broadly categorized as (a)Feature detection and classification using hand-crafted techniques (b)Combination of hand-crafted and machine learning techniques (c)Employing Deep learning techniques especially CNN’s for feature extraction and classification.

One of the earliest automated systems using handcrafted technique given by Siddalingaswamy and Gopalakrishna [14] used clustering and mathematical morphological techniques to detect exudates. In [15] the authors adopted marker-controlled watershed transformation for extracting exudates to perform DME stage classification. The work given in [16] used top-down image segmentation and local thresholding to find the region of interest, followed by exudate detection. Deepak and Jayanthi [11] used supervised learning approach to capture the global characteristics in fundus images and assessed disease severity using a rotational asymmetric metric by examining the symmetry of the macular region. The work proposed in [17] used a combination of handcrafted and machine learning techniques. This work detected macula by locating darkest pixel along enhanced blood vessels followed by forming clusters of these pixels where the largest cluster formed was considered as the macula. They used Gabor filter as well as adaptive thresholding to detect exudates, and grade DME severity with the help of support vector machine (SVM). The work given in [18] proposed a method for texture extraction from various regions of interest of the fundus image and employed SVM for grading severity of DME.

However, the overall performance of the above-described grading systems is largely based on feature extraction strategies, exudate segmentation and the location of anatomical structures i.e. macula and fovea. In addition, the extraction of features is hugely dependent on the dataset being used to evaluate the methodology. Finding and developing a feature set that is suitable for different datasets remains a challenge, as features that describe one dataset may not often describe other datasets. Thus deep learning emerged as a more promising approach to learn features automatically. One of the most recent works [19] used CNN’s to automatically extract features and grade the input fundus images. The authors obtained an accuracy of 88.8%, sensitivity of 74.7% and specificity of 96.5%.

One of the earliest works in the domain of ensemble learning was proposed by Dasarthy and Sheela [20] which deals with partitioning of feature space using two or more classifiers. Recently ensemble learning techniques applied on computer vision problems in the domain of medical imaging with CNN’s as classifiers showed improvement in the performance [21, 22]. Feature space expansion and use of different starting points has been instrumental in providing better approximation of the optimal solution and has been cited as the rationale for the improved classification using ensemble learning. Ensemble learning offers promising alternative to the currently used techniques in computer vision domain and we wish to explore it in our proposed DMENet algorithm.

## Material and Preparation

### Datasets

The proposed model was trained using data from two publicly available datasets, IDRiD [23] and MESSIDOR [24]. Both these datasets are graded on a scale of three, where each grade is described as given in Table 1.

**Table 1.** Description of grades

| Grade | Description |
| --- | --- |
| 0 | No Apparent hard Exudate(s) |
| 1 | Presence of hard Exudate(s) outside the radius of one disc diameter from the macula center |
| 2 | Presence of hard Exudate(s) within the radius of one disc diameter from the macula center |

### IDRID

This database contains 516 fundus images. These images were captured by a retinal specialist at an Eye Clinic located in Nanded, Maharashtra, India. Experts verified that all images are of adequate quality, are clinically relevant, no image is duplicated and ensured that a reasonable mixture of disease stratification representative of DME is present. The images have a resolution of 4288×2848 pixels. The dataset contains 222 image of grade 0, 41 of grade 1 and 243 images of grade 2. Images were acquired using a Kowa VX-10 alpha digital fundus camera with a 50-degree field of view (FOV), and all are centered near to the macula.

### MESSIDOR

The Messidor database has been established to facilitate studies on computer-assisted diagnoses of diabetic retinopathy. The database contains 1200 fundus images acquired at three different locations. The images have resolutions of 1440*960, 2240*1488 and 2304*1536 pixels. This dataset contains 974, 75 and 151 images of grade 0, 1 and 2 respectively. Images were acquired using a color video 3CCD camera on a Topcon TRC NW6 non-mydriatic fundus camera with a 45-degree field of view.

### Data Preparation

#### Data Preprocessing

Preprocessing plays a key role in improving accuracy of results by reducing noise in the background. It removes some of the variations between images due to differing lighting conditions and camera resolution thus making the data consistent [25]. The images contained in the above-mentioned databases have different image resolutions were scaled down to a fixed resolution size to form a standardized dataset and also to reduce computational costs. The standardized dataset obtained was preprocessed to improve image contrast which would help in clear distinction of exudates from the background. The preprocessed image is obtained in a two-step process where we first perform image enhancement and follow it up with the appplication of the morphological opening operation on the output of the first step. The enhancement of the input image follows the mathematical formula as given in equation 1.

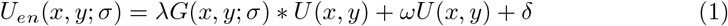

where *U_en_* denotes the corresponding output image after enhancement, *U*(*x, y*) denotes the raw fundus image, *G*(*x, y*; *σ*) is a Gaussian kernel with scale *σ* [25]. The convolution operator is represented by *. In this paper *σ* is empirically set as 25, *λ*, *ω* and *δ* are parameters to control the weights, which are empirically set as −4, 4 and 0.5.

We then perform morphological opening on the enhanced image to control the brightness of blood vessels which appear brighter than other retinal surfaces due to the lower reflectance [26]. It performs the second step using the expression given by equation 2.

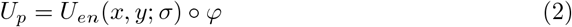

where *U_p_* denotes the corresponding output image after performing morphological opening and is the final output of preprocessing stage. *ϕ* represents the structuring element disk with a radius of 3 pixels and o denotes a morphological opening operation. Fig. 1 shows comparision of the preprocessed image with respect to the original image.

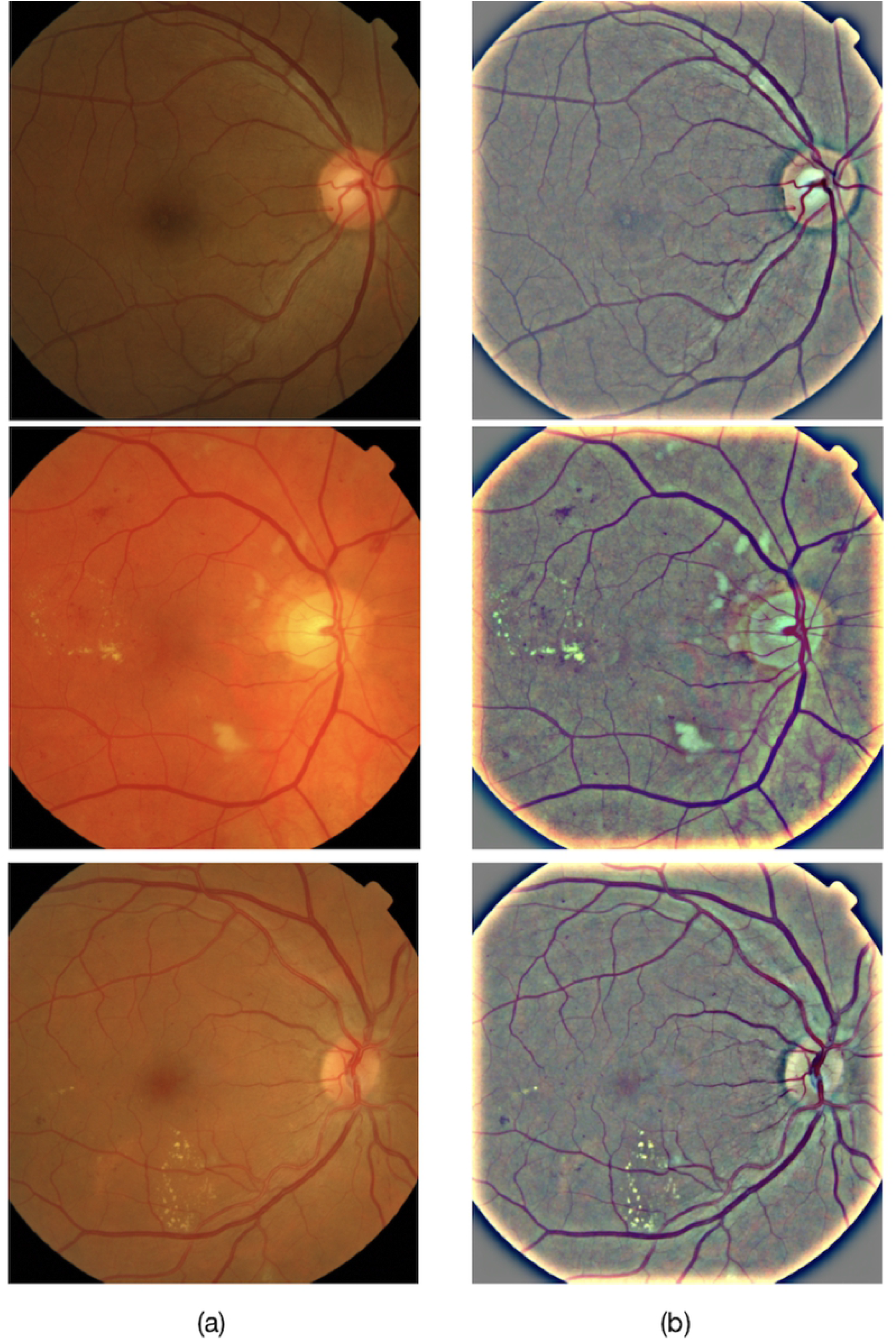

#### Data Augmentation

In order to build a robust automated disease grading system using CNN’s it is important to have a dataset of images with uniform representation from all the classes. In the field of Medical imaging, datasets are often imbalanced, as the number of patients that turn out to be positive is very less compared to total cases. This class imbalance problem introduces significant challenge when training deep CNN’s which are data intensive [26]. Another common issue that occurs while training deep CNN’s on smaller datasets is overfitting [27]. Training deep CNN’s on larger datasets has shown to improve robustness and generalisability of the model [25]. Data augmentation is an effective solution to reduce overfitting during CNN training as well as to balance the samples across different classes. There are different augmentation techniques like Flips, Gaussian-Noise, Jittering, Scaling, Gaussian-Blur, Rotations, Shears, etc which are widely used. Flips, Rotations, and Scaling have outperformed other techniques [4]. Our augmentation methods include random rotations between 0 to 40 and horizontal flips.

### Proposed Methodology-DMENet

In this section, we present the details of the proposed methodology-DMENet. It is a two-stage pipeline that performs disease classification (as Positive DME and Negative DME) in the first stage followed by severity grading (as Grade 1 and Grade 2) in the second stage. Only those images that are classified as positive DME are passed onto the second stage. The main workflow of our algorithm is illustrated in Fig. 2. Each stage comprises of a classification structure (referred to as Gamma ensemble in first stage and Delta ensemble in second stage) based on our proposed novel ensemble technique known as *Hierarchical Ensemble of Convolutional Neural Networks* (HE-CNN). To delve deeper into the ensemble methodology it is prudent to understand CNN’s architecture and finetuning technique which are described below.

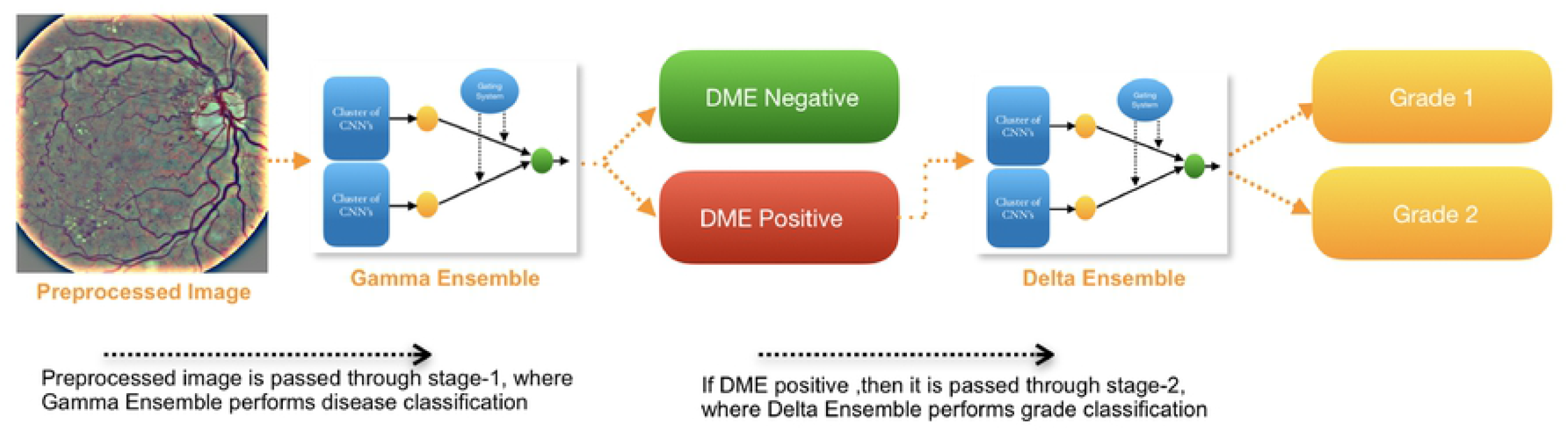

#### Background: CNN and Finetuning

It is important to have a brief understanding of the Convolutional Neural Networks (CNN’s) which forms the pillar of the proposed solution. CNN’s are used in biomedical image analysis since 1990’s [28, 29], however, with the advent of GPU’s and availability of larger and better datasets, they have started showing superior performance. The strength of CNN lies in its deep architecture which is responsible for extracting distinguishing features at different layers of abstraction [30, 31]. CNN’s are fundamentally made of three types of layers, namely convolutional, pooling, and fully-connected layers. The convolutional layer is composed of a set of convolutional kernels which are responsible to learn the patterns or specific features from the input. These kernels compute different feature maps and each neuron in a feature map is associated with a region of neighboring neurons of the previous layer. We can obtain the new feature map by convolving the input with a trained kernel. We then apply element-wise nonlinear activation function on the results obtained using convolution operator [32]. Activation functions are really important for CNN’s to learn and understand non-linear complex functional mappings between the inputs and the response variable. Consider 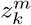 to be the *m^th^* feature map (output) of layer *m*. The feature map *k* for *m^th^* convolutional layer can be mathematically represented as follows

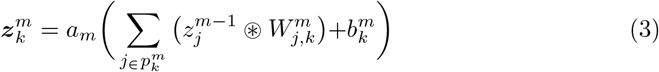

where *a_m_*(.) denotes the activation function, 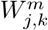 is the weight vector of the filter that connects feature map *j* in layer *m* − 1 to feature map *k* in layer *m* and 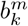 is bias term associated with *k^th^* feature map in the *m^th^* layer. 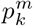 denotes the set of planes in layer *m* − 1 that are connected to *k^th^* feature map. Suppose the size of input feature map 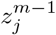 is *L*^*m*−1^ × *B*^*m*−1^ pixels, and size of convolutional masks 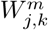 is *c^m^* × *f^m^* pixels. The Fig. 3 illustrates the convolutional layer. The size of output feature map would be accordingly (*L*^*m*−1^ − *c*^*m*^ − 1) × (*B*^*m*−1^ − *f^m^* − 1) pixels.

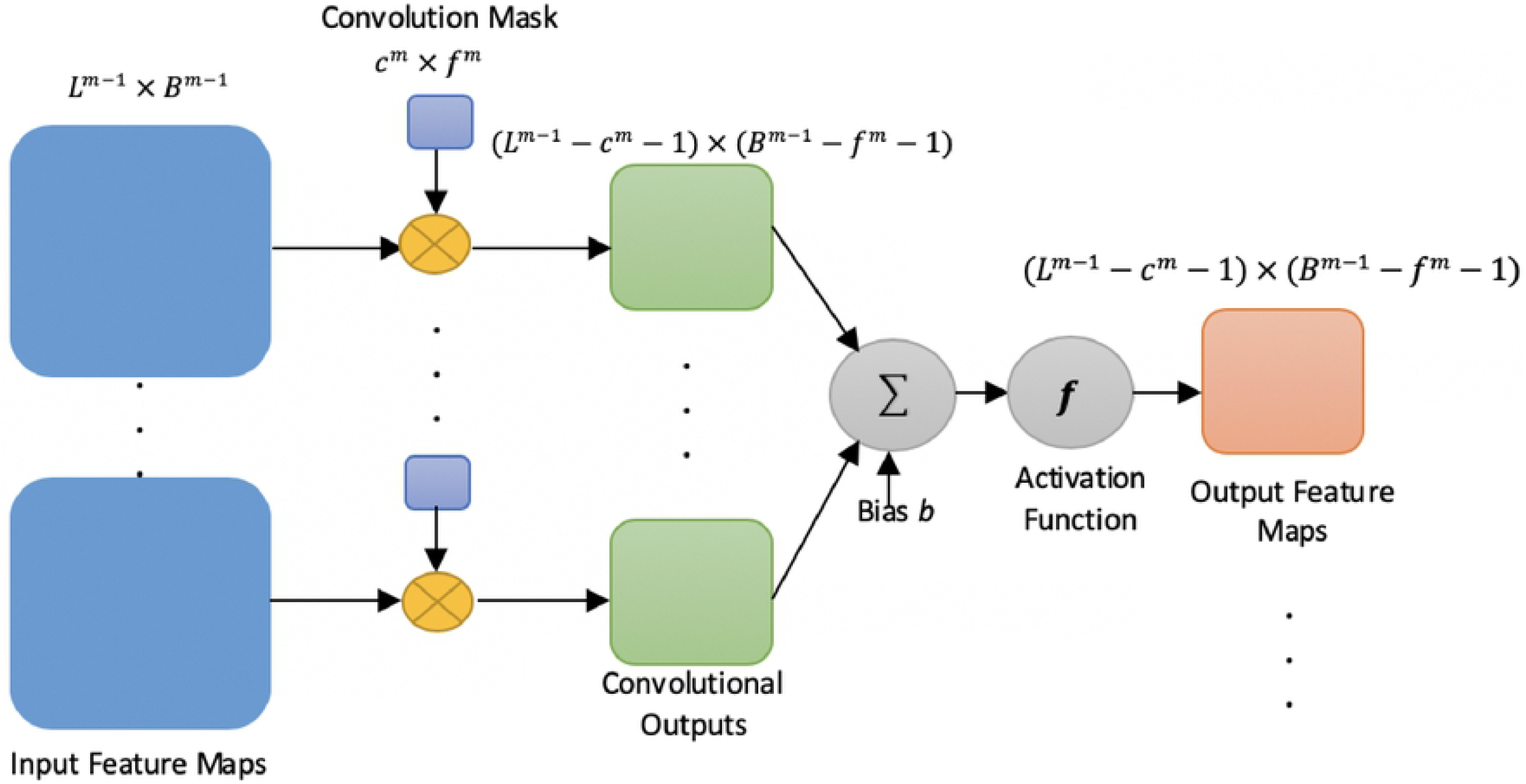

The pooling layer is placed between two convolutional layers with the aim of reducing spatial dimensions, improving the computing performance and reducing the number of parameters that control overfitting. There are various types of pooling operations however the most commonly used ones are max-pooling [33], and average pooling [34]. The feature map *k* for *m^th^* pooling layer can be mathematically represented as follows

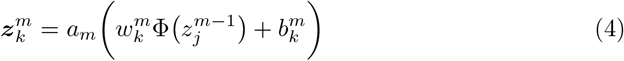

where Φ(.) denotes the pooling function, 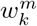 and 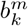 are the scalar weights and the bias term respectively. Here we divide the feature map *k* of convolution layer (*m* − 1) into non-overlapping blocks of size 2 × 2 pixels. In case of max-pooling we take the largest element from the rectified feature map within that block. We slide our 2 × 2 block by 2 cells (also called ‘stride’) and take the maximum value in each region. Clearly, a feature map 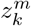 in pooling layer m will have a size of *L^m^* × *B^m^* where *L^m^* = *L*^*m*−1^/2 and *B^m^* = *B*^*m*−1^/2. The Fig. 4 shows the pooling layer.

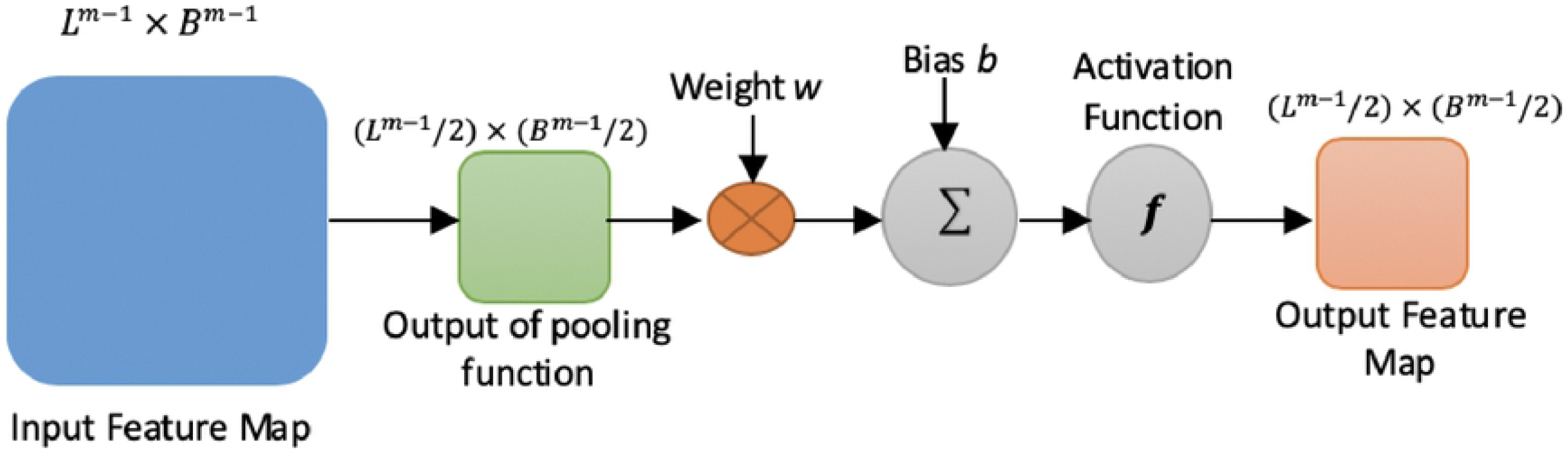

After several convolutional and pooling layers, we have a fully connected layer which performs high-level reasoning [35, 36]. A fully connected layer takes all the neurons in the previous layer and connects them to each of the current layer’s single neurons to generate global semantic information. [32]. The result of the final output layer is given as

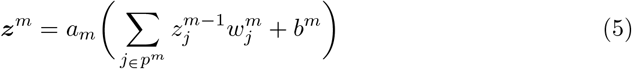

Various studies [37, 38] demonstrated that transfer learning or finetuning a CNN is better than training a CNN from scratch. Training CNN’s from scratch suffers from several issues especially the requirement of having a huge dataset is a serious concern in the medical domain where the expert annotation is a costly affair. Training a CNN from scratch is also computationally expensive, time-consuming and suffers from frequent overfitting and convergence issues. An effective alternative for training a CNN from scratch is transfer learning. Transfer learning is the method where CNN’s extract generic features from smaller datasets based on its feature learning from a larger and well-labeled dataset. The factors which affect the transfer learning strategy is the size of the new dataset on which it is applied and its similarity to the original dataset [39]. CNN’s learn low-level features in the early layers which are general for any network whereas high level features are learnt in later layers that are dataset dependent. One of the more advanced approach of transfer learning is Fine-tuning which employs the back-propagation algorithm to update the pre-trained weights *w* of a CNN’s. Fine-tuning is an iterative process that works by minimizing the cost function with respect to the pre-trained weights [22]. The cost function is represented as follows

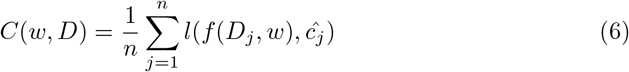

where *D* is training dataset with n images, *d_j_* is the *j^th^* image of *D, f* (*d_j_, w*) is the CNN function that predicts the class *c_j_* of *d_j_* given *w,ĉ_j_* is the ground-truth of *j^th^* image, *l*(*c_j_, ĉ_j_*) is a penalty function based on logistic loss *l* for predicting *c_j_* instead of *ĉ_j_*.

In order to minimize the cost function, the Stochastic Gradient Descent algorithm [40] is commonly used where the cost over entire training dataset is calculated based on the approximated results obtained over mini-batches of data.

If 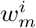 denotes the weights in the *m^th^* convolutional layer at iteration *i*, and *Ĉ* denotes the cost over a mini-batch of size y, then the updated weights in the next iteration are computed as follows [38]

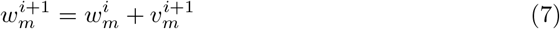

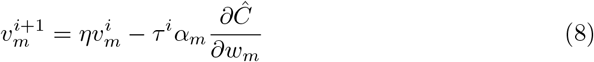

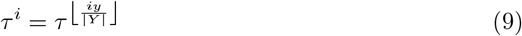

where |*Y*| denotes the number of training images, *α_m_* is the learning rate of the *m^th^* layer which controls the size of updates to the weights, *η* is the momentum coefficient that indicates the contribution of the previous weight update in the current iteration. This has the effect of speeding up the learning process while simultaneously smoothing the weight updates, *τ* is the scheduling rate that decreases learning rate *α* at the end of each epoch.

#### Proposed Classification Methodology-Hierarchical Ensemble of Convolutional Neural Networks

This section introduces our novel classification structure known as Hierarchical Ensemble of Convolutional Neural Networks (HE-CNN) which is used to design the Delta and Gamma ensembles present in the DMENet pipeline as shown in Fig. 2. An overview of the HE-CNN is shown in Fig. 5 (We have shown HE-CNN representation of a two-level architecture, however, this methodology can be generalized for n levels). The proposed HE-CNN architecture comprises multiple learners that maps the input to the output and contains various Gating Systems that characterize the hierarchical structure of the architecture. There is a Local Gating System (LGS) for each cluster of learners and a Global Gating System (GGS) that serves to consolidate the outputs of these clusters. The output of the *i^th^* cluster is given by

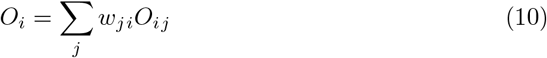

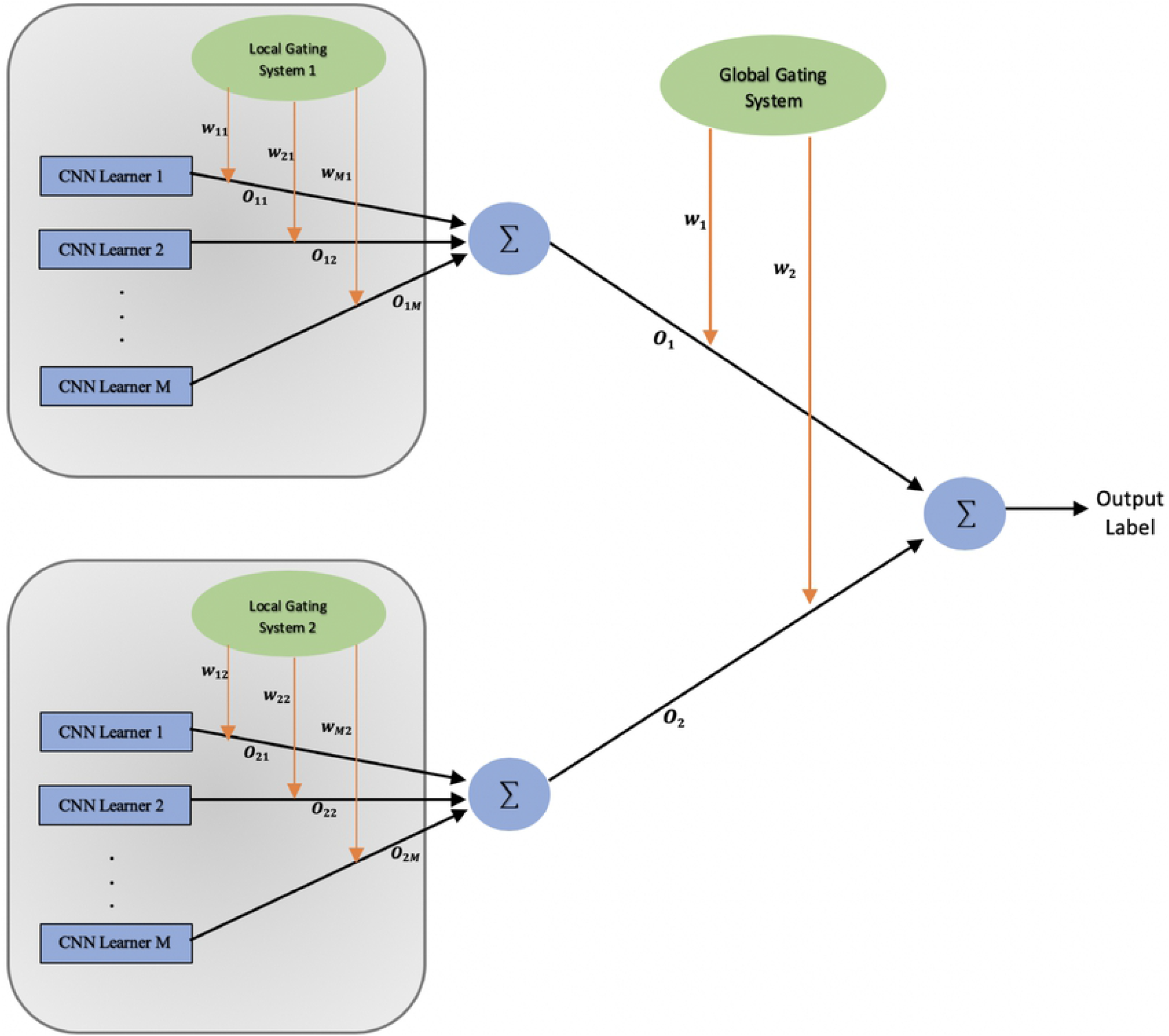

where *w_ji_* is activation of *j^th^* output unit of Local Gating System in the *i^th^* cluster 2. and *O_ij_* denotes the output given by *j^th^* learner in the *i^th^* cluster.The final output of 2. the HE-CNN architecture is given by 2

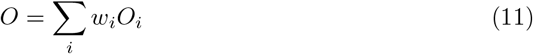

where *w_i_* is activation of *i^th^* output unit of Global Gating System and *O_i_* denotes the output given by *i^th^* cluster.The outputs of the gating systems are normalised using softmax function [41]

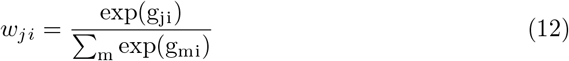

and

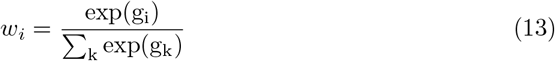

where *g_i_* and *g_ji_* are the weighted sums at the output units of the corresponding gating networks.

As shown in Fig. 5, in this architecture we are dividing the input space into a nested set of regions and then attempting to fit simple surfaces to the data which fall in these regions. The regions would be having soft boundaries which means that the data points are spread across multiple regions and the boundaries between these regions are simple parameterized surfaces which are adjusted from the learning process [42]. Every learner has expertise in one specific area of the high-dimensional input space and each learner estimates the conditional posterior probability on the partitioned feature space separated by a gating system based on the given input. This model combines the outputs of several CNN Learners by training the gating system. The gating systems in the architecture are basically classifiers responsible for dividing the input space. Their decision of division is based on the learners capability to model the input-output functions within their respective regions as quantified by their posterior probabilities. We have analysed the hierarchical nature of this architecture using probabilistic interpretation as given below. We then discuss the error functions that were employed for training simultaneously both the CNN learners and the gating systems.

##### Probabilistic Interpretation

The HE-CNN model learns from each cluster which have multiple CNN learners that are experts in their respective regions. Each *f_j|i_* (here *i* denotes the cluster and *j* denotes the learner in *i^th^* cluster) maps the input x to one of the output classes *G* i.e. in Gamma Phase-positive or negative DME and in Delta Phase-Grade 1 or Grade 2. Each *j^th^* learner in *i^th^* cluster maps the input x to one of the output classes *G* (In Gamma Phase-DME or No DME and in Delta Phase-Grade 1 or Grade 2). The total probability of generating *G* from x is the mixture of the probabilities of generating *G* from each of the component densities *P*(*G, x*; *O_ij_*), where the mixture components are multinomial probabilities:

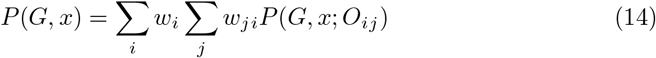

Here *P*(.) represents the likelihood function of the HE-CNN model. The gating system is responsible for assigning weights and thereby allowing the overall model to execute a competitive learning process by maximizing the likelihood function of the training data [43–45]. Each learner module specializes in exclusive regions of feature space and all modules in this methodology learn simultaneously by interacting with each other rather than learning independently [46]. The learners in a cluster are closely linked with each other and learn similar mappings early in the training phase. They differentiate later in training as the probabilities associated with the cluster to which they belong become larger. Thus the architecture tends to acquire coarse structure before acquiring fine structure, this particular feature of the architecture is notable as it provides robustness to problems with overfitting in deep hierarchies.

##### Error Function

Taking the error functions used in [42, 44] we have made suitable changes to adapt to our model. Assuming a training set of *V* images, where the error function of CNN learners for *v^th^* input image *x_v_* is defined as:

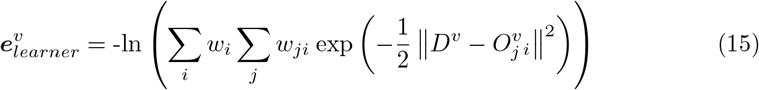

where *D^v^* is the desired output vector, 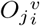 is the output vector of *j^th^* learner in *i^th^* cluster. The effective error for *j*^th^ learner in *i*^th^ cluster on full training set is defined as

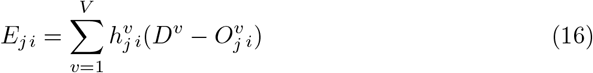

where 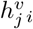 is the posterior probability estimate provided by *j^th^* learner of *i^th^* cluster for input *x_v_* and 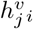 is given as

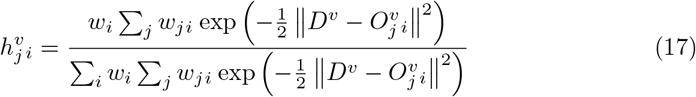

The error functions of both the gating systems (GGS, LGS) are defined as follows

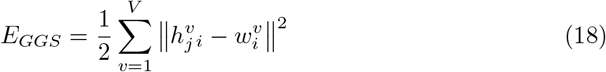

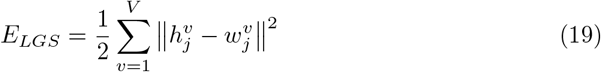

where 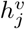 gives the posterior probability provided by *j^th^* learner in the cluster for which error function is being calculated. It is defined as given below.

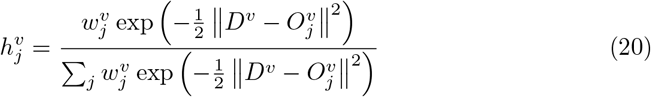

## Experiments, Results and Discussion

### Evaluation Metrics and Experimental Strategy

To assess the performance of our proposed DMENet methodology and HE-CNN ensemble we have employed standard performance measures. The Confusion Matrix and Receiver Operating Characteristic (ROC) analyses were used to calculate the accuracy, precision, sensitivity/recall, specificity and average area under the ROC curves (AUC). F1–score is the harmonic mean of precision and recall [47]. F1–score is calculated as given below.

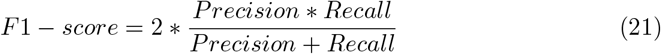

F1–score is a more robust metric to evaluate the classification performance as it takes into consideration the class imbalance problem by giving equal importance to precision and recall thus involving both false positives and false negatives. F1–score ranges between 0 and 1, reaches the best value 1 when the balance between precision and recall is perfect. Cohens kappa (*κ*) score is used to determine the potential of HE-CNN in partitioning the feature space [48]. *κ* measure signifies the level of classifiers agreement and its value ranges between −1 and 1 (higher the *κ* score higher is the agreement).

In this study, we have applied 5-fold cross-validation technique to assess the classification measures and generalize the performance of the model. The dataset was roughly split into five equal-size partitions and ensured that each partition has good representation as a whole. By preserving random seed across all iterations, the dataset-dependent bias is eliminated. Four partitions were used for training and the rest of the partition were used for testing. This step has been repeated five times until all the different test set choices have been evaluated. Over the five folds, the classification measures were averaged.

### Experiment Design

This study is based on a two-stage DMENet methodology which uses HE-CNN as a key classification algorithm in each stage. One of the main issue we faced while developing the DME screening solution is the class-imbalance problem as the datasets which we are using contain a significant portion of DME negative images and more importantly the grade 1 images are less than 10% of the whole dataset. This led to mis-classification of an image with grade 1 characteristics, thus we have broken the tri-class (Grade 0, 1, 2) problem into a binary classification problem using two stage DMENet model as depicted in Fig. 2. We have used a combination of pre-trained networks like ResNet [49], DenseNet [50], SqueezeNet [51], GoogleNet [31] and SE-ResNet [52] as the learners in each cluster of the ensemble model. The gating systems, both the local and global are based on the CNN architectures given in Table 2. The final implementation details of each classification structure (i.e. Gamma and Delta Ensemble) after performing number of experiment’s are given below.

**Table 2.** CNN Structures

| Structure | Input Size | Number of layers | First Convolution mask | Other Convolutional masks | Max-Pooling Size | Stride | Pool type in FC | Number of output neurons |
| --- | --- | --- | --- | --- | --- | --- | --- | --- |
| CNN-1 | 224 x 224 | 18 | 5 x 5 | 3 x 3 | 2 x 2 | 2 | Average | 2 |
| CNN-2 | 224 x 224 | 22 | 5 x 5 | 3 x 3 | 2 x 2 | 2 | Average | 2 |
| CNN-3 | 224 x 224 | 28 | 7 x 7 | 3 x 3 | 2 x 2 | 2 | Average | 2 |
| CNN-4 | 224 x 224 | 32 | 7 x 7 | 3 x 3 | 2 x 2 | 2 | Average | 2 |
| CNN-5 | 224 x 224 | 40 | 7 x 7 | 3 x 3 | 2 x 2 | 2 | Average | 2 |

*Gamma Ensemble* follows the *two-level* HE-CNN architecture as shown in Fig. 5. The CNN learners in the first cluster are setup using a combination of ResNet and DenseNet architectures. We used a pretrained ResNet-50 as learner-1, pretrained DenseNet-161 as learner-2 and the outputs of these learners were aggregated using LGS whose architecture is based on the CNN-1 structure as given in Table 2. The second cluster is composed of three CNN learners based on SqueezeNet, SE-ResNet and ResNet architectures. We used pretrained SE-ResNet-50 as learner-1, pretrained SqueezeNet as learner-2 and pretrained ResNet-34 architecture as learner-3. The CNN-2 structure given in Table 2 is used as LGS for aggregating the outputs. In order to combine the outputs of both these clusters, we used a GGS which used CNN-3 architecture as given in Table 2.

*Delta Ensemble* is based on *three-level* HE-CNN architecture. It contains an additional cluster of CNN learners and corresponding LGS as compared to the two-level architecture of Gamma Ensemble. The first cluster contains two pretrained CNN learners which are based on the architectures of DenseNet and SE-ResNet. We used DenseNet-169 as learner-1, SE-ResNet-50 as learner 2 and we combined the outputs of these learners using an LGS-1 which used CNN-2 architecture as given in Table 2. The two pretrained CNN learners in the second cluster are based on architectures of ResNet-50 and SqueezeNet. We used a CNN-1 architecture given in Table 2 as LGS-2 to combine the outputs of the learners in the second cluster. The third cluster is composed of three pretrained CNN learners-DenseNet-161, ResNet-34 and GoogLeNet. The CNN-3 architecture is employed as LGS-3 and the CNN-4 architecture as given in Table 2 for GGS.

All the CNN learners were initialized on the ImageNet [53] dataset weights. The filter weights derived from ImageNet were then finetuned through back-propagation thus minimizing the CNN’s empirical cost in equation 6. We used Stochastic Gradient Descent as discussed in section to minimize the cost function. Here the cost calculated over mini-batches of size 32 is used to approximate the cost over the entire training set. We set the learning rate *α_m_* to 10^-3^ ensuring proper convergence and the scheduling rate which depends on the speed of convergence *τ* is set to 0.9. The training process for both the gating systems LGS and GGS is performed using Adam optimizer [54] with the learning rate of 10^-2^, batch size of 32, *β*_1_ = 0.9, *β*_2_ = 0.999 and decay of 10^-5^.

### Results and Discussion

In this experiment, we performed analysis of various sorts ranging from evaluating the proposed HE-CNN ensemble technique in each stage of DMENet pipeline, comparative evaluation of the DMENet methodology with existing computer-aided solutions, analyzing performance of CNN’s v/s proposed HE-CNN technique, comparative study of HE-CNN with other existing ensemble techniques and finally analyzing the performance of DMENet v/s tri-class classification (Grade 0, Grade 1 and Grade2). All the analysis and comparisons are made using the evaluation metrics as discussed in the section.

#### Evaluation of Gamma and Delta Ensembles

We demonstrate the performance in each stage of DMENet pipeline using Confusion Matrix and ROC curves. The Confusion Matrix in Fig. 6.A shows the performance of Gamma Ensemble on 360 test images and Fig. 6.B gives the performance of Delta Ensemble on 244 test images. Table 3 shows the results obtained by Gamma and Delta ensembles on various metrics of accuracy, specificity,sensitivity, precision, F1–score and Kappa score. The ROC curves of the Gamma and Delta Ensembles are shown in Fig. 7. The AUC scores obtained by the Gamma and Delta Ensembles are 0.9654 and 0.9489 respectively. We can clearly see that the Gamma and Delta ensembles achieved promising results.

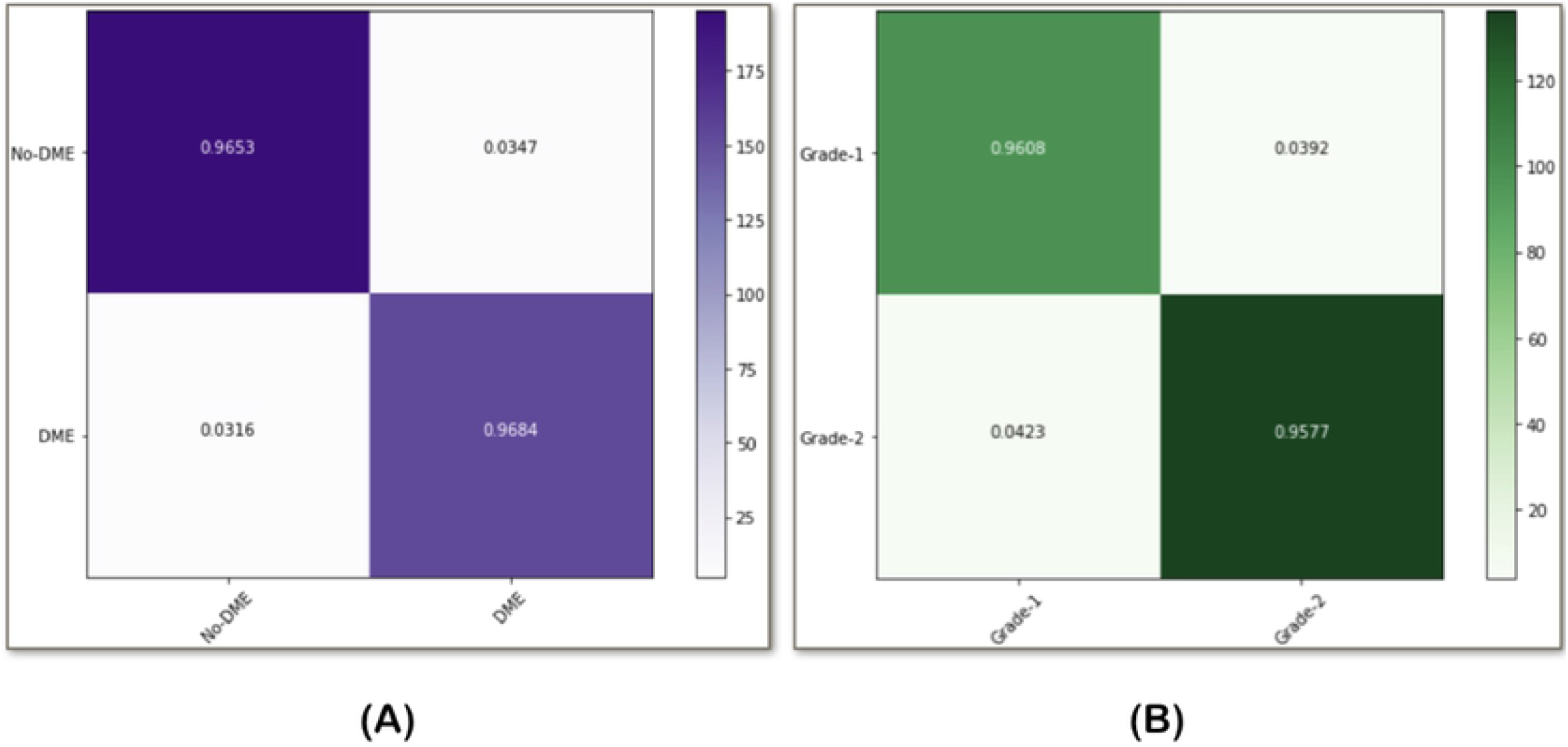

**Table 3.** Performance of Gamma and Delta Ensembles

|  | Gamma Ensemble | Delta Ensemble |
| --- | --- | --- |
| *Accuracy* | 96.67 | 95.91 |
| *Specificity* | 95.63 | 97.14 |
| *Sensitivity* | 97.50 | 94.23 |
| *Precision* | 96.53 | 96.08 |
| *F1-score* | 0.9701 | 0.9515 |
| *Kappa-Score* | 0.932 | 0.916 |

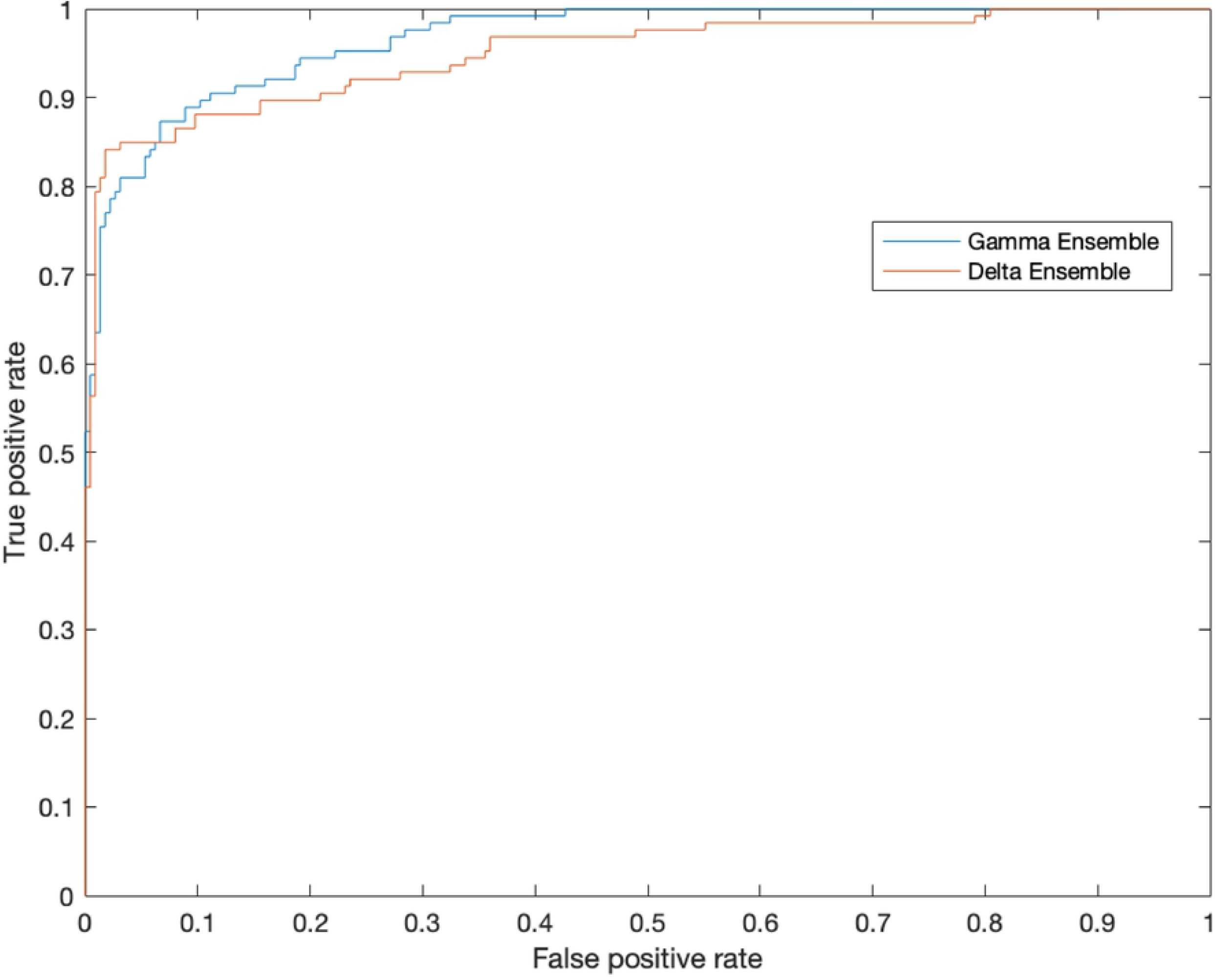

#### Comparative Evaluation of HE-CNN Methodology

To obtain a benchmark for comparing the performance of HE-CNN ensemble and to show its capability, we are evaluating our model with regard to non-ensemble methods using independent CNN’s as well as some of the existing ensemble techniques on both the classification tasks in DMENet methodology. We have analyzed the performance of CNN’s using pretrained ResNet, DenseNet architectures and the reults obtained are shown in Table 4. However, we can see from the results that these models did not perform well as they were overfitting the given data. Also prevential measures were taken to control overfitting by adding dropout factor of 50% on the fully connected layer before the output. Although dropout helped in reducing overfitting but the results obtained are not satisfactory as compared to results given by HE-CNN ensemble. To elaborate the comparative performance of HE-CNN we used some standard ensemble techniques as follows. Ave-Ensemble averages the output maps of independently trained CNN’s to generate the final output [55, 56]. Soft-Ensemble employes several parallel CNN branches where feature maps are concatenated at the end of each convolutional branch and fully-connected layer processes all the features [57], [58]. Mixture-Ensemble uses multiple CNN’s and a weight regulation network [59]. Pruned-Ensemble uses pruning to retain best pretrained models and then make predictions using maximum voting [60]. The results of these ensemble techniques can be observed in Table 4. All the above models are evaluated independently on two different classification tasks as in DMENet methodology (i.e. DME and severity classification), the results obtained are then mapped to a tri-class problem in order to measure various metrics. This is done in order to understand the true power of HE-CNN architecture. Exact numerical values of DMENet’s performance in comparison to other existing methodologies is shown in Table 7.

**Table 4.** Comparitive study of HE-CNN ensemble

| Classifier | Performance Measures |  |  |  |
| --- | --- | --- | --- | --- |
|  | Accuracy | Specificity | Sensitivity | F1-score |
| *CNN Models* |  |  |  |  |
| ResNet-34 | 82.16 | 94.27 | 70.97 | 0.7923 |
| ResNet-34+Dropout | 83.65 | 91.38 | 74.49 | 0.8216 |
| DenseNet-169 | 74.21 | 92.26 | 58.35 | 0.7175 |
| DenseNet-169+Dropout | 78.43 | 87.34 | 64.83 | 0.7345 |
| *Ensemble Models* |  |  |  |  |
| Ave-Ensemble | 83.62 | 86.42 | 78.27 | 0.8132 |
| Soft-Ensemble | 84.56 | 82.84 | 87.39 | 0.8421 |
| Pruned-Ensemble | 88.78 | 91.23 | 88.14 | 0.8967 |
| Mixture-Ensemble | 93.16 | 92.39 | 93.94 | 0.9274 |
| HE-CNN | 96.12 | 95.84 | 96.32 | 0.9609 |

In the HE-CNN architecture, one of the key component was the gating systems (both LGS and GGS) and we have used adam optimizer while training them. We compared SGD with Nesterov momentum (with the learning rate set as 10^-2^ and momentum set as 0.9) to adam optimization method and came to a conclusion that adam optimizer performed well in this task. The results obtained are shown below in Table 5.

**Table 5.**
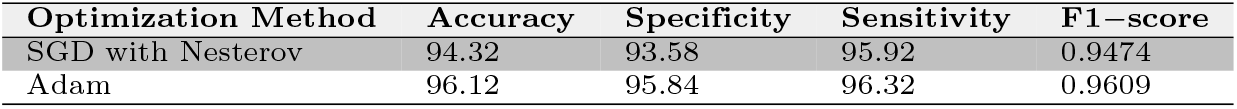
Results of Optimization Methods

| Optimization Method | Accuracy | Specificity | Sensitivity | F1-score |
| --- | --- | --- | --- | --- |
| SGD with Nesterov | 94.32 | 93.58 | 95.92 | 0.9474 |
| Adam | 96.12 | 95.84 | 96.32 | 0.9609 |

We have analyzed the performance of DMENet two-stage pipeline with respect to tri-class problem to show the competence of DMENet pipeline in DME screening. The fundus images used in DME screening are generally graded into one of the three classes (Grade 0, 1 and 2). The results obtained by performing tri-class classification using pretrained CNN’s as well as ensemble methods using HE-CNN are shown in Table 6 along with results obtained using the corresponding pretrained CNN and HE-CNN. The results strongly validates our premise of breaking the tri classification problem into binary classification using two levels.

**Table 6.** Results of DMENet v/s Tri-class classification

| Optimization Method | Accuracy | Specificity | Sensitivity | F1-score |
| --- | --- | --- | --- | --- |
| ResNet-34<br>(Tri-Class) | 68.31 | 83.28 | 67.21 | 0.7114 |
| ResNet-34<br>(DMENet) | 82.16 | 94.27 | 70.97 | 0.7923 |
| HE-CNN<br>(Tri-Class) | 91.46 | 88.84 | 89.62 | 0.8849 |
| HE-CNN<br>(DMENet) | 96.12 | 95.84 | 96.32 | 0.9609 |

**Table 7.** Comparitive study of DMENet’s performance with recent solutions for DME screening

| Author | Year | Technique | Database | Results |
| --- | --- | --- | --- | --- |
| Siddalingaswamy<br>*et al.* [14] | 2010 | Clustering and morphological techniques | 148 images acquired from the Department of ophthalmology, Kasturba medical college, Manipal (Local Dataset) | Sensitivity-95.6%<br>Specificity-96.15 |
| Lim *et al.* [15] | 2011 | Marker- controlled watershed transformation for extracting exudates and performed DME stage classification using the location of extracted exudates | MESSIDOR- Publicly Available Database | Sensitivity-80.9%<br>Specificity-90.2%<br>Accuracy-85.2% |
| Jaafar *et al.* [16] | 2011 | Based on top-down image segmentation and local thresholding by a combination of edge detection and region growing Grading of hard exudates was performed using a polar coordinate system centred at the fovea | DIARETDB0 ,DIARETDB1 and MESSIDOR- Publicly Available Databases | Sensitivity-93.2%<br>Specificity-90.5% |
| Akram *et al.* [17] | 2012 | Filter Bank and SVM classifier | MESSIDOR- Publicly Available Database | Sensitivity-92.6%<br>Specificity-97.8%<br>Accuracy-97.3% |
| Aditya *et al.* [18] | 2015 | Texture extraction and SVM classifier | Local Dataset | Sensitivity-91%<br>Specificity-75%<br>Accuracy-86% |
| Baidaa *et al.* [19] | 2016 | Convolutional Neural Networks | MESSIDOR- Publicly Available Database | Sensitivity-74.7%<br>Specificity-96.5%<br>Accuracy-88.8% |
| Proposed DMENet | 2019 | Hierarchical Ensemble of CNN’s (HE-CNN) | MESSIDOR, IDRiD- Publicly Available Database | Sensitivity-96.32%<br>Specificity-95.84%<br>Accuracy-96.12% |

## Conclusion

In this paper we have successfully demonstrated the efficacy of our proposed DMENet methodology and HE-CNN architecture. The proposed methodology is simple yet extremely efficient. We strongly believe that the HE-CNN architecture is fairly general and with little modification, it has the potential of solving all such classification problems in the field of bio-medical imaging elegantly.

